# Binocular modulation of monocular V1 neurons

**DOI:** 10.1101/320218

**Authors:** Kacie Dougherty, Michele A. Cox, Jacob A. Westerberg, Alexander Maier

**Affiliations:** Department of Psychology, College of Arts and Science, Vanderbilt Vision Research Center, Center for Integrative and Cognitive Neuroscience, Vanderbilt University, Nashville, TN 37203, USA

**Keywords:** binocular vision, monocular neurons, binocular combination, binocular summation, binocular integration

## Abstract

In humans and other primates, sensory signals from each eye remain separated until they arrive in the primary visual cortex (V1), but their exact meeting point is unknown. In V1, some neurons are activated by stimulation of only one eye (monocular neurons) while most neurons are driven by stimulation of either eye (binocular neurons). Monocular neurons are most prevalent in the main input layers of V1 while binocular neurons dominate the layers above and below. This observation has given rise to the idea that the two eyes’ signals remain separate until they converge outside V1’s input layers. Here, we show that despite responding to only one eye, monocular neurons in all layers, including the input layers, of V1 discriminate between stimulation of their driving eye alone and stimulation of both eyes. This finding suggests that binocular signals occur at an earlier processing stage than previously appreciated as even so-called monocular neurons across all V1 layers encode what is shown to both eyes.

## INTRODUCTION

Front-facing eyes are one of the most prominent features that differentiate humans and other primates from their closest ancestors (Heesy, 2004). As a consequence, our eyes’ views overlap, which forces our brains to combine the outputs of the eyes to yield a singular view (Blake et al., 1981, Harwerth and Smith, 1985, Ohzawa and Freeman, 1986, Sengpiel and Blakemore, 1996, Ohzawa et al., 1997a, Ohzawa et al., 1997b, Livingstone and Tsao, 1999, Cumming and DeAngelis, 2001, Parker and Cumming, 2001, Ding and Sperling, 2006, Meese et al., 2006, Parker, 2007, Freeman, 2017). For this binocular combination to occur, the signals from the eyes need to meet at some point along the primary visual pathway. While the possible locations underlying this convergence have been narrowed, the exact meeting point of the two eyes’ signals is unknown.

Primate retinae do not receive feedback from the visual structures to which they project (Reperant et al., 1989). This connectivity suggests that the output of each eye remains entirely separate from that of the other. Following visual transduction in the retina, the monocular signals from retinal ganglion cells mainly project to the lateral geniculate nucleus of the thalamus (LGN) (Casagrande and Boyd, 1996). For almost all primate LGN neurons, visually stimulating one eye leads to a response (dominant eye) while stimulating the other does not (non-dominant, or silent eye) (Wiesel and Hubel, 1966). In other words, stimulation of each eye separately evokes responses in two mutually exclusive groups of LGN neurons. It thus seems that the formation of a binocular signal occurs at a subsequent stage of visual processing.

As a next step in the primary visual pathway, afferents from the LGN mainly project to the primary visual cortex (V1) (Casagrande and Boyd, 1996). LGN neurons that respond to stimulation of the same eye innervate the same neurons in V1 layer 4 (termed layer 4C in primates) (Hubel and Wiesel, 1972). In line with this connectivity, many layer 4C neurons, like their LGN counterparts, do not respond when a stimulus is shown to one of the eyes (Hubel and Wiesel, 1968, Blasdel and Fitzpatrick, 1984). A popular interpretation of these findings is that the signals from each eye remain largely segregated in layer 4C of V1, with binocular convergence happening at a subsequent step of processing.

Layer 4C neurons converge onto neurons in the layers above (Mitzdorf, 1985, Douglas, 1989). The prevailing model of binocular convergence builds on the findings listed above by proposing that V1 neurons in superficial layers of V1 receive inputs from layer 4C neurons that respond to one eye as well as from layer 4C neurons that respond to the other eye (Hubel and Wiesel, 1972). Indeed, most supragranular neurons respond to either eye (albeit to varying degrees) (Hubel and Wiesel, 1965, Hubel and Wiesel, 1968, Hubel and Wiesel, 1972). Neurons in the supragranular layers project to neurons in the lower layers of V1, which also respond to stimulation of either eye (Casagrande and Boyd, 1996).

One challenge to the model outlined above is that layer 4C neurons also receive inputs from cortical neurons in addition to the inputs from the LGN (Ahmed et al., 1994, Binzegger et al., 2004). These intracortical connections raise the interesting possibility that even monocular layer 4C neurons that are driven by one eye exclusively encode a binocular signal. This seemingly paradoxical situation could arise if the firing rates of monocular neurons change reliably when both eyes are stimulated simultaneously. In other words, even though stimuli shown to their silent, non-dominant eye alone do not evoke responses, monocular neurons might nonetheless systematically modulate responses when stimuli are shown to both eyes. Indeed, some monocular V1 neurons are tuned for interocular disparity, which demonstrates that they are sensitive to what is shown to both eyes (Ohzawa and Freeman, 1986, Prince et al., 2002, Read and Cumming, 2004). However, whether such neurons can also be found in layer 4C is unclear because most of the previous studies lacked laminar resolution.

Here we use laminar neurophysiology to determine whether the signals from the two eyes truly remain segregated in V1 layer 4C. To do so, we examined the extent to which neurons in all layers of V1 are sensitive to one or both eyes. Specifically, we employed linear multielectrode arrays to record V1 laminar neural responses in macaques that viewed stimuli with one eye, the other eye or both eyes simultaneously. We found that 78% of V1 neurons across all V1 layers were binocular in that they were significantly driven when stimuli were presented to either eye. In line with earlier work, we located the bulk of monocular neurons to layer 4C. Strikingly, we found that, although activated by only one eye, these so-called monocular neurons responded significantly differently when both eyes were stimulated simultaneously. This phenomenon of binocular modulation occurs across all layers of V1, and affects both orientation tuned and non-orientation tuned monocular neurons in layer 4C. These findings suggest that, despite their name, monocular neurons in the primary input layers of V1 actually encode both eyes’ views. Thus, binocular signals are present at an earlier processing stage than commonly thought.

## RESULTS

In each session, we penetrated the *dura mater* over V1 with a linear multielectrode array and positioned the array so that its contacts spanned the depth of cortex (Maier et al., 2010, Cox et al., 2017, Dougherty et al., 2017). While we recorded extracellular voltages, we displayed visual stimuli through a mirror stereoscope to stimulate the eyes independently (**Figure 1a**). After mapping the population receptive field (RF) location for the neurons under study (see **STAR METHODS**), we presented static sine-wave gratings to the left eye, right eye, or both eyes over the RF location (**Figure 1b**). Offline, we extracted spiking activity from recorded extracellular voltage data (see **STAR METHODS**).

**Figure 1.**
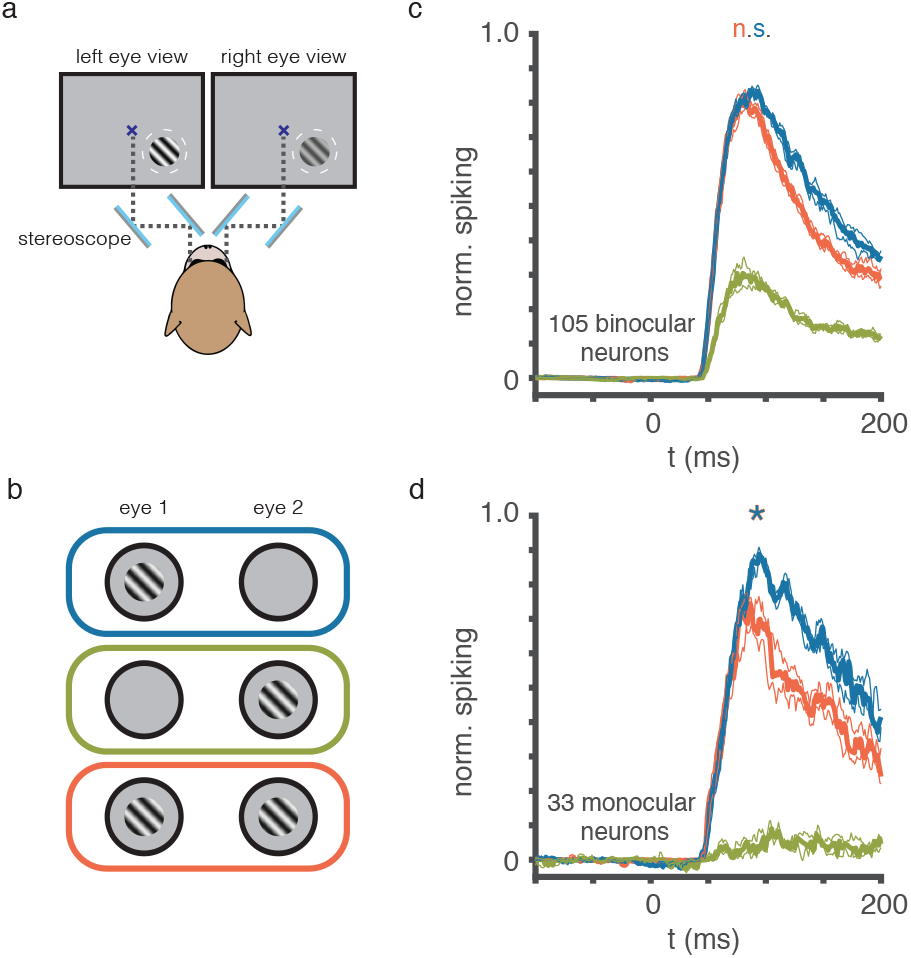
The Majority but Not All V1 Neurons Are Driven Through Both Eyes. (a) Top-down view of experimental setup. Visual stimuli were presented to fixating macaque monkeys through a mirror stereoscope that consisted of two pairs of mirrors. The mirrors were angled in a way that each eye of the animal saw the left and right halves of the monitor, respectively. (b) On every stimulus presentation, a sine-wave grating was presented over the RF location to either the left eye (blue), right eye (green), or both eyes (orange). (c) Median normalized spiking responses of binocular V1 neurons to the dominant eye (blue), both eyes (orange), or the non-dominant eye (green). There was no significant difference between the response to the dominant eye and the response to both eyes (two-tailed Wilcoxon signed-rank test, p = 0.092, N = 105). Thin lines around each median represent 75% confidence intervals (chosen to account for the fact that the data was not normally distributed). (d) Median normalized responses of monocular V1 neurons to the same stimulation conditions as in (c). All conventions as in (c). Note that stimulating the dominant eye in isolation evoked a significantly larger population spiking response than stimulating both eyes simultaneously (twotailed Wilcoxon signed-rank, p = 0.029, N = 33). Response to the non-dominant eye was not significant at α = 0.05. *See also Figure S1*.

### The Vast Majority of V1 Neurons Are Driven Through Both Eyes

We collected visual responses for 290 neurons throughout all V1 layers across both animals (261 from E48, 29 from I34). Congruent with previous work (Hubel and Wiesel, 1968, Blasdel and Fitzpatrick, 1984), monocular stimulation of either eye led to a statistically significant response (paired t-test, α = 0.05) for the majority of neurons (n = 226) (orange and green traces in **Figure 1c**). Monocular neurons that responded to only one eye (paired t-test, α = 0.05) made up just 22% of the population (n = 64) (**Figure 1d**).

### Binocular Modulation among Binocular V1 Neurons

For a subset of neurons in our sample (n = 138), we collected responses to zero disparity, matching stimulation of both eyes (dioptic) simultaneously. Binocular neurons, that responded to stimulation of either eye, exhibited a small, non-significant response difference between dominant eye and binocular stimulation (n = 105, two-tailed Wilcoxon signed-rank test, p = 0.65) (**Figure 1c**). In other words, on the population level, binocular V1 neurons respond similarly when their dominant eye is stimulated in isolation or when both eyes are stimulated simultaneously. This finding is congruent with previous work (Smith et al., 1997, Truchard et al., 2000), suggesting that binocular V1 responses are normalized to account for the increase in sensory input when both eyes are stimulated rather than one. However, note that close to half of the individual binocular V1 neurons in our sample either significantly increased or decreased their firing rates when both eyes were stimulated rather than one (**Table 1**, Wilcoxon rank sum test, p < 0.05).

**Table 1.** Fraction of binocular neurons with significant binocular response modulation,

|  | *two-tailed rank sum*<br>*test* | *left-tailed rank sum*<br>*test (facilitated)* | *right-tailed rank*<br>*sum test*<br>*(suppressed)* |
| --- | --- | --- | --- |
| *binocular neurons,*<br>*$p < 0.05$* | 40/105 | 28/105 | 23/105 |

### Responses of V1 Monocular Neurons Modulate During Binocular Stimulation

We repeated the same analysis for our population of monocular neurons. If monocular neurons are truly sensitive to only one eye, there should be no measurable difference between monocular and binocular stimulation. Instead, we found that the population response of monocular neurons showed a significant difference between binocular and monocular stimulation (n = 33, two-tailed Wilcoxon signed-rank test, p = 0.029) (**Figure 1d**, see also **Figure S1a**). Fixational eye movements did not affect this result (**Figure S1a**).

We further investigated this binocular modulation by subdividing the monocular neurons into two groups: The first group, *binocularly suppressed neurons*, included all monocular neurons whose firing rates decreased during binocular stimulation. The second group, *binocularly facilitated neurons*, included all neurons whose firing rates increased during binocular stimulation. Using this categorization scheme, the suppressed group included over two-thirds of all monocular neurons (n = 23, one-tailed Wilcoxon signed-rank test, p =1.44 × 10^−5^) (**Figure 2a**). The facilitated group included the other third of monocular neurons (n = 10, one-tailed Wilcoxon signed-rank, p =9.77 × 10^−4^) (**Figure 2b**).

**Figure 2.**
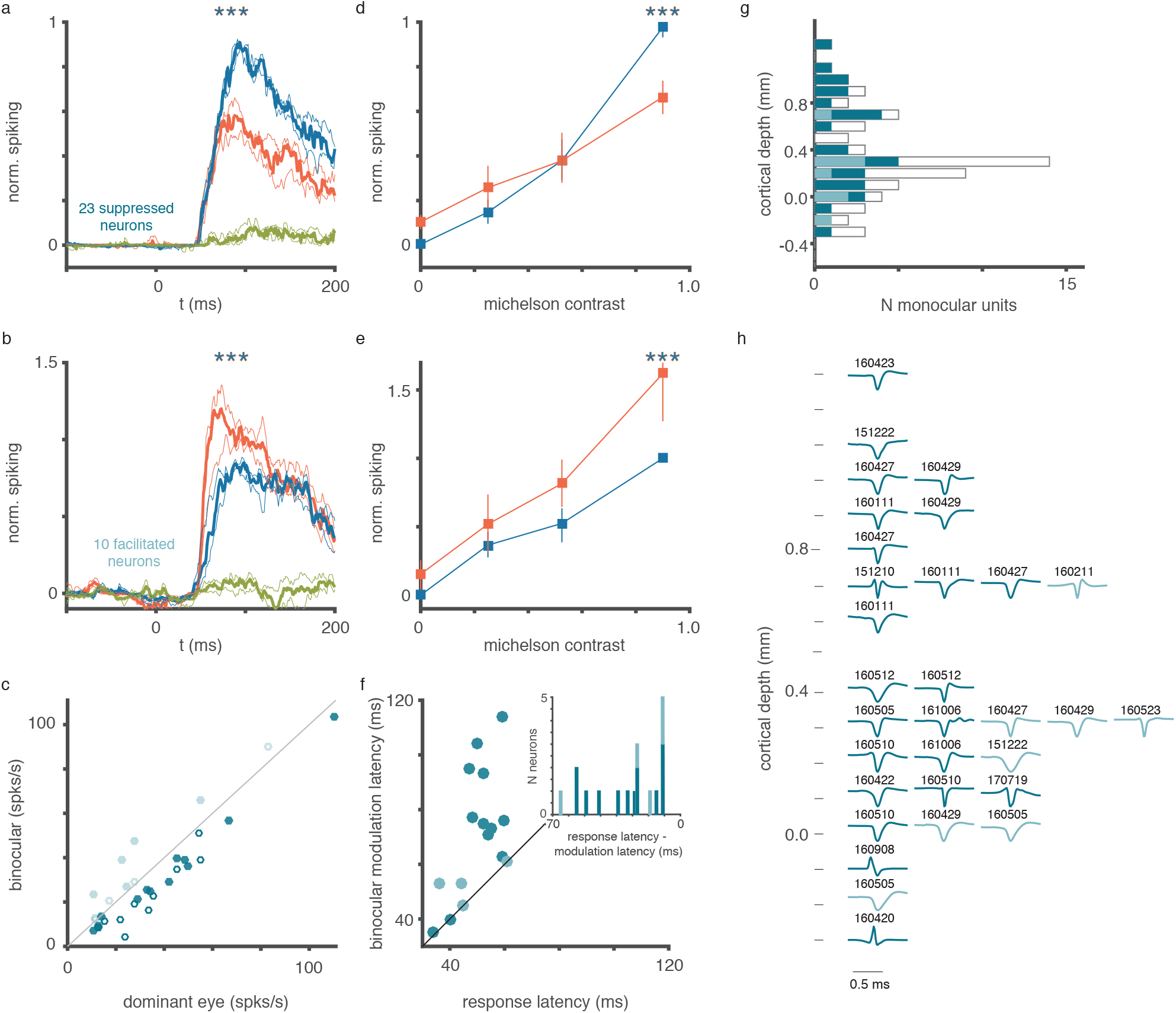
Responses of V1 Monocular Neurons Modulate During Binocular Stimulation. (a) Median normalized responses of monocular neurons that reduced firing rates during binocular stimulation. Their population response to binocular stimulation differed significantly from monocular stimulation (one-tailed Wilcoxon signed-rank, p = 1.44 x 10^−5^, n = 23). All conventions as in Figure 1c,d. Response to the non-dominant eye was not significant at α = 0.05. (b) Same as (a) but for monocular neurons that increased spiking during binocular stimulation. Their population response under binocular stimulation was significantly greater than for stimulating their dominant eye alone (one-tailed Wilcoxon signed-rank, p = 9.77 x 10^−4^, n = 10). (c) Mean dominant eye response versus mean binocular response for each monocular neuron. In all cases, spiking averages were computed across the initial response period (40-140 ms). Solid circles indicate neurons with a difference in firing rate at α = 0.1 (see **STAR METHODS**). (d) Mean normalized contrast responses for suppressed monocular neurons during dominant eye stimulation (blue, non-dominant eye 0.0 contrast) and binocular stimulation (orange, 0.8 or greater contrast). All contrast values and sample sizes are detailed in Table 1. Error bars represent 95% confidence limits. When the contrast of the stimulus in the dominant eye equal was 0.8 or greater, neurons were significantly suppressed (one-tailed, Wilcoxon signed-rank test, p = 2.15 × 10^−5^, Bonferroni corrected for multiple comparisons). Differences at all other contrast levels were not significant. (e) Same as (d) but for facilitated monocular neurons. Binocular stimulation resulted in significantly greater spiking responses than stimulating dominant eye alone (one-tailed, Wilcoxon signed-rank test, p = 9.77 × 10^−4^, Bonferroni corrected for multiple comparisons). (f) Response latency versus onset of binocular modulation for neurons with significant binocular modulation. Conventions as in (c). Modulation latency corresponds to the first time point at or after the response latency with a response difference at α = 0.1. Histogram depicts the time difference between response latency and binocular modulation latency for suppressed and facilitated monocular neurons. (g) Laminar sites for all monocular neurons across the cortical depth (0 mm corresponds to the CSD-determined L4C/L5 boundary, see **STAR METHODS**). White bars indicate the location of all monocular neurons (including those for which we were unable to record binocular data). Dark green bars indicate suppressed monocular neurons and light green bars indicate facilitated monocular neurons. (h) Average spike waveforms for all suppressed and facilitated monocular neurons at their relative position across cortical depth. Colors as in (c), (f), (g). *See also Figures S1 and S3*.

We repeated this comparison for a more liberally-defined group of monocular neurons (one-tailed t-test, p > 0.01). Using this liberal criterion yielded a larger sample size (n = 51) at the expense of including neurons that showed a minimal response to the non-dominant eye. Nonetheless, this liberally-defined group yielded comparable results to those shown in **Figure 2a** and **2b** (**Figure S1d**, one-tailed Wilcoxon signed-rank, p = 6.17 × 10^−7^, N = 31; **f**, one-tailed Wilcoxon signed-rank, p = 4.78 × 10^−5^, N = 20).

Lastly, we compared the firing rate of each individual V1 unit under binocular and monocular stimulation (**Figure 2c**). Congruent with the group statistics outlined above, most neurons showed significant facilitation or suppression (solid symbols in **Figure 2c**; see **STAR Methods**).

### Binocular Modulation Occurs at High Contrasts and Early in the Response

All analyses so far were performed using stimuli of high visual contrast (see **STAR Methods**). Psychophysical studies have shown that binocular combination differs at varying contrast levels (Legge and Rubin, 1981, Ding and Sperling, 2006). We therefore wondered if stimulus contrast affects the binocular modulation of monocular neurons demonstrated above. To test for the impact of stimulus contrast, we recorded the responses of a subsample of monocular neurons to binocular stimuli presented at several contrast levels (see **Table 1, Table S1** for N). Specifically, we either did not display a stimulus (monocular conditions) or showed a high contrast grating (binocular conditions) to the non-dominant eye. We then paired these conditions with stimuli of varying contrast in the dominant eye (**Table 1**).

For the suppressed group, binocular modulation only occurred when a high contrast stimulus was present in the dominant eye (one-tailed, Wilcoxon signed-rank test, p = 2.15 x 10^−5^, Bonferroni corrected for multiple comparisons) (**Figure 2d**). Likewise, facilitated neurons only exhibited binocular modulation whenever we presented a high contrast stimulus in the dominant eye (one-tailed, Wilcoxon signed-rank test, p = 9.77 x 10^−4^, Bonferroni corrected for multiple comparisons) (**Figure 2e**). Repeating this analysis for the liberally-defined group of monocular neurons yielded a similar result (see **Figure S1e,g Table S1**). These results suggest that monocular V1 neurons modulate their firing rates at relatively high visual contrast levels only, mirroring the psychophysical observation that binocular combination differs fundamentally between high and low contrast levels.

We next wanted to know whether binocular modulation occurred early or late in the visual response. Such timing differences are informative because more sophisticated intracortical processing, involving neurons in several layers, likely takes more time than direct interactions within layer 4C. Accordingly, we considered the onset latency of binocular modulation across our population of monocular neurons (see **STAR Methods**). This analysis revealed significant binocular modulation at the onset of the visual response for neurons in both the facilitated and suppressed groups. Overall, the onset of binocular modulation occurred somewhat earlier for facilitated neurons (median: 53 ms) compared to suppressed neurons (median: 75 ms) (**Figure 2f**).

### Binocular Modulation Occurs in the Primary Input Layer of V1

The prevailing model of binocular processing suggests that signals from each eye remain separate in the retino-geniculate input layer 4C of V1 before merging to a binocular response in the layers above (Hubel and Wiesel, 1972). If most or all the monocular neurons we recorded were located outside layer 4C, our results fit the model. If, on the other hand, monocular neurons in layer 4C showed similar sensitivity to both eyes, the model would need to be revisited. To test between these two possibilities, we next strove to locate binocularly modulating monocular neurons within the laminar microcircuit of V1.

We used current source density analysis (CSD; see **STAR Methods**) to estimate the location of each neuron relative to the layer 4C-layer 5 boundary. Congruent with previous work (Hubel and Wiesel, 1968, Hubel and Wiesel, 1972), we found that the majority of monocular neurons were located in layer 4C (defined as 0.0 to 0.5 mm relative to L4C/L5 boundary), with a smaller fraction of monocular neurons in the layers above and below (**Figure 2g, Figure S1i**). Most facilitated monocular neurons were located in granular layer 4C. In contrast, suppressed monocular neurons were evenly distributed between supragranular and granular layers (**Figure 2g,h**).

We were able to collect responses to monocular and binocular stimuli for 16 monocular neurons in the extragranular layers outside of L4C and 17 neurons inside granular layer 4C. Seven of the extragranular monocular neurons (43%), and eight of the monocular L4C neurons (47%) exhibited a significant response difference between dominant eye and binocular stimulation (ROC analysis, α = 0.05) (**Figure S1b**, see **STAR Methods**). However, the population response of the nine neurons that did not significantly modulate when both eyes were stimulated, showed a similar trend (**Figure S1c**).

We were able to determine the orientation tuning for five L4C monocular neurons. Three of these neurons showed a significant effect for orientation (ANOVA, p < 0.05). Of those three neurons, two neurons also showed a significant effect of binocular modulation (ROC analysis, α = 0.05). Of the two untuned, monocular L4C neurons, one showed significant binocular modulation (**Figure S3a**). Thus, even some untuned monocular L4C neurons are sensitive to both eyes. We also obtained orientation tuning data for seven more monocular neurons outside of L4C. All of these neurons showed a significant orientation tuning effect (ANOVA, p < 0.05).

Importantly, the waveforms of these monocular neurons did not resemble the tri-phasic waveforms associated with axonal spikes (Lemon and Prochazka, 1984), suggesting that we did not mistake LGN afferents for V1 neurons (**Figure 2h**). Furthermore, the characteristics of monocular neurons in our sample resembled those of previous reports: First, most binocular neurons (99%) were tuned for orientation, whereas almost half of monocular neurons (45%) were not (**Figure S3b**) (Hubel and Wiesel, 1968, Hubel and Wiesel, 1977, Blasdel and Fitzpatrick, 1984). Second, the baseline firing rates of the monocular neurons were overall significantly higher than that of binocular neurons (two-sample t-test, t_288_ = 4.83, p = 2.21 x10^−6^) (**Figure S3c**) (Snodderly and Gur, 1995 but see Blasdel and Fitzpatrick, 1984).

### Ocular Dominance Varies Across V1 Layers

How do the above findings relate to the well-documented V1 phenomenon of ocular dominance (Hubel and Wiesel, 1968, Hubel and Wiesel, 1977), which describes neuronal preferences for one eye over the other eye? To answer this question, we estimated ocular dominance by computing an ocularity index for all single neurons and multiunits (see **STAR METHODS**), and then compared relative ocular dominance across cortical depth (see also Hubel and Wiesel, 1968, Schiller et al., 1976, Hubel and Wiesel, 1977). This ocularity index was defined as the Michelson contrast between the responses to monocular stimulation of each eye. Accordingly, a value of −1 corresponds to neurons exclusively driven through the ipsilateral eye, a value of 1 corresponds to neurons driven exclusively through the contralateral eye, and a value of 0 corresponds to neurons driven equally through either eye.

Our sample spanned the entire index range of ocular dominance (**Figure 3a**). Across the neuronal population, the spread of ocular dominance resembled a normal distribution (Chi-square goodness of fit, χ^2^ = 9.64, d.f. = 7, p = 0.21) (**Figure 3b**). We repeated this analysis for multiunits (see **STAR Methods**), as these data provided a larger sample. Multiunit responses reflect the activity of neurons up to 350 μm away (Mineault et al., 2013). This distance can bridge neighboring ocular dominance columns (Wiesel et al., 1974, Florence and Kaas, 1992, Horton and Hocking, 1996). Given this shortcoming, we expected multiunits to exhibit a stronger bias towards binocular responses. Indeed, the mean rectified ocularity indices for multiunits were lower than those for single neurons (one-tailed t-test, p = 1.98 x 10^−25^, *t*_1285_ = 10.58). Similar to the single neuron population, multiunit ocular dominance was normally distributed (Chi-square goodness of fit, χ^2^ =3.95, d.f. = 7, p = 0.79) (**Figure 3c**), suggesting that the mode of V1 neurons respond with equal magnitude to either eye.

**Figure 3.**
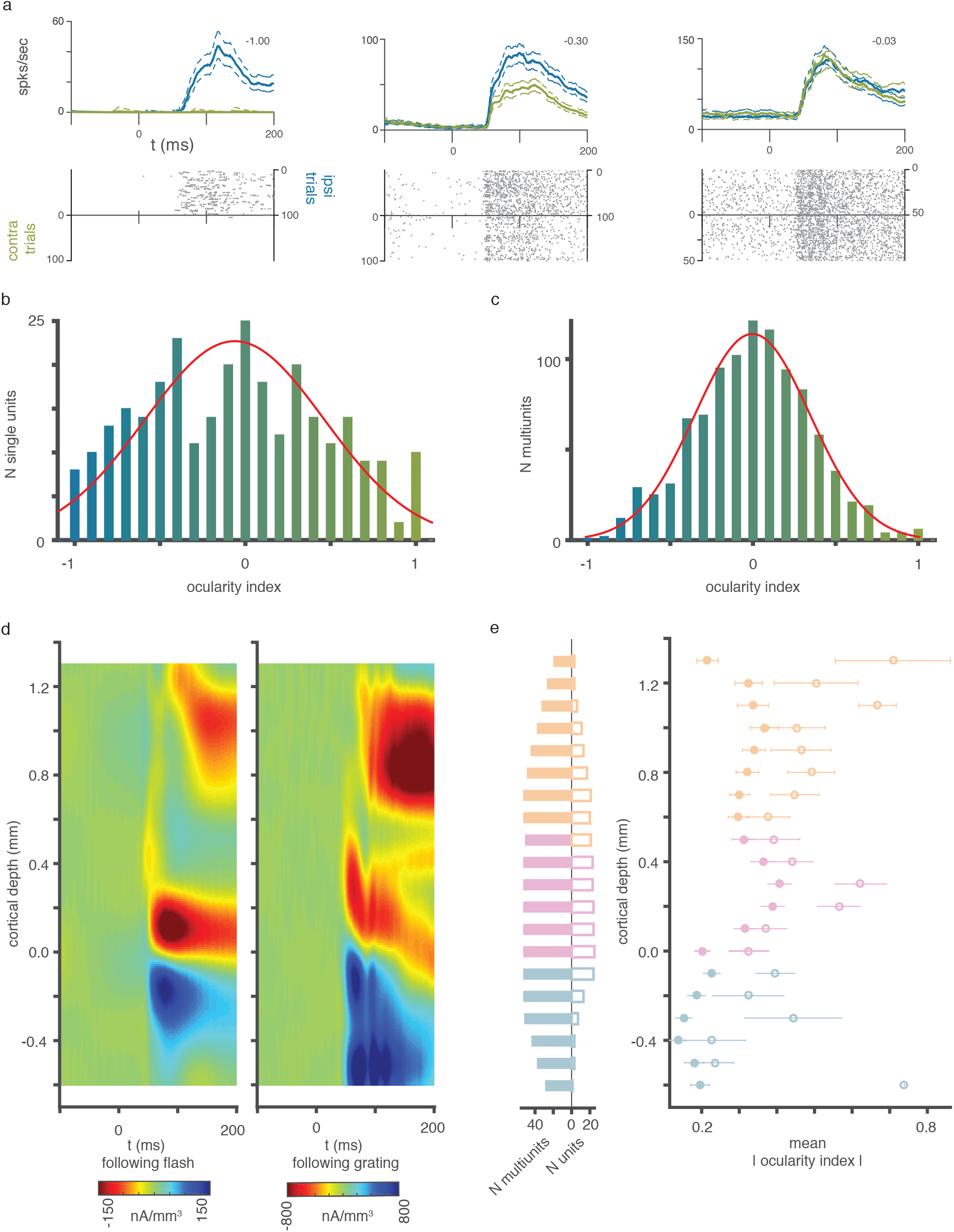
Ocular Dominance Across V1 Layers. (a) Three example neurons of varying ocular dominance. Spike density functions for their relative responses to the non-dominant eye are shown in green. Responses to their dominant eye are shown in blue. Corresponding raster plots are shown below. The resulting ocularity index (see **STAR METHODS**) for each neuron is shown in top right corner. Thin, dashed lines represent 95% confidence intervals around the mean response. (b) Histogram overlaid with fitted Gaussian (red) for the ocularity indices of all recorded single neurons (n = 290, Chi-square goodness of fit, χ^2^ = 9.64, d.f. = 7, p = 0.21). (c) Same as (b) but for all multiunits recorded in our sample (n = 997). Red fit is Gaussian (Chi-square goodness of fit, χ^2^ =3.95, d.f. = 7, p = 0.79). (d) Left panel: Mean CSD response to a full-field white flash (N = 33 penetrations, both animals). Right panel: Mean CSD response to a sine-wave grating presented at the RF location (N = 45 penetrations, both animals). 0 mm marks the L4C/L5 border, estimated using the bottom of the initial current sink (see **STAR METHODS**). (e) Number of multiunits (solid symbols) and number of single neurons (hollow symbols) (left) included in the computation of the mean rectified ocularity index at each cortical depth (right). *See also Figure S4*.

We next determined the profile of ocular dominance across the cortical depth of V1, using the boundary between layer 4C and layer 5 as our reference point (0.0 mm) (**Figure 3d**, see also **Figure S4**). We then calculated the mean rectified ocularity index of all neurons at each cortical depth and used these values as a measure of how much each neuron was driven through one or both eyes (**Figure 3e**). Note that using absolute values means that an index of 0 corresponds to equal responses to each eye.

Previous work demonstrated greater ocular dominance among layer 4C neurons (Hubel and Wiesel, 1968, Blasdel and Fitzpatrick, 1984). This observation can be explained by the fact that (monocular) geniculate inputs primarily target this layer (Casagrande and Boyd, 1996). Indeed, we found that, on average, both single neurons and multiunits exhibited their strongest preference for one eye over another in layer 4C. For the multiunits in particular, the rectified ocularity indices differed significantly between laminar compartments (one-way ANOVA, *F*(2,981) = 62.51, p = 2.80 x 10^−26^), with a significant difference between upper and lower layers (Tukey’s HSD post-hoc test, p = 9.56 x 10^−10^, N_supragranular_ = 342, N_infragranular_ = 326) as well as between middle and lower layers (Tukey’s HSD post-hoc test, p = 9.56 x 10^−10^, N_granular_ = 316, N_infragranular_ = 326). The ocularity in upper and middle layers was not significantly different (Tukey’s HSD post-hoc test, p = 0.47, N_supragranular_ = 342, N_granular_ = 316).

Overall the pattern of rectified ocularity indices across cortical depth was qualitatively comparable for multiunits and single neurons, with some divergence at the upper and lower bounds that might be explained by the lower number of single neurons sampled at these locations (**Figure 3e**).

### Ocular Dominance Correlates with Binocular Modulation

As described above, we observed a wide range of ocular dominance among our sample of V1 neurons. We also found that the majority of V1 neurons are binocular, and that their combined response does not show any significant binocular modulation. In contrast, binocular modulation could be observed for both our conservatively- and liberally-defined populations of monocular neurons. These observations led us to ask if there was a systematic relationship between ocular dominance and binocular modulation.

To test for this relationship, we calculated a binocular modulation index that quantifies both the strength and direction of binocular modulation (see **STAR METHODS**). We then assessed whether a neuron’s ocular dominance had any explanatory power for that neuron’s binocular modulation. Interestingly and largely congruent with the findings above, we found a significant correlation between the binocular modulation index and the rectified ocularity index (p = 0.0011, r^2^ = 0.0756, r = −0.275) (**Figure 4a**).

**Figure 4.**
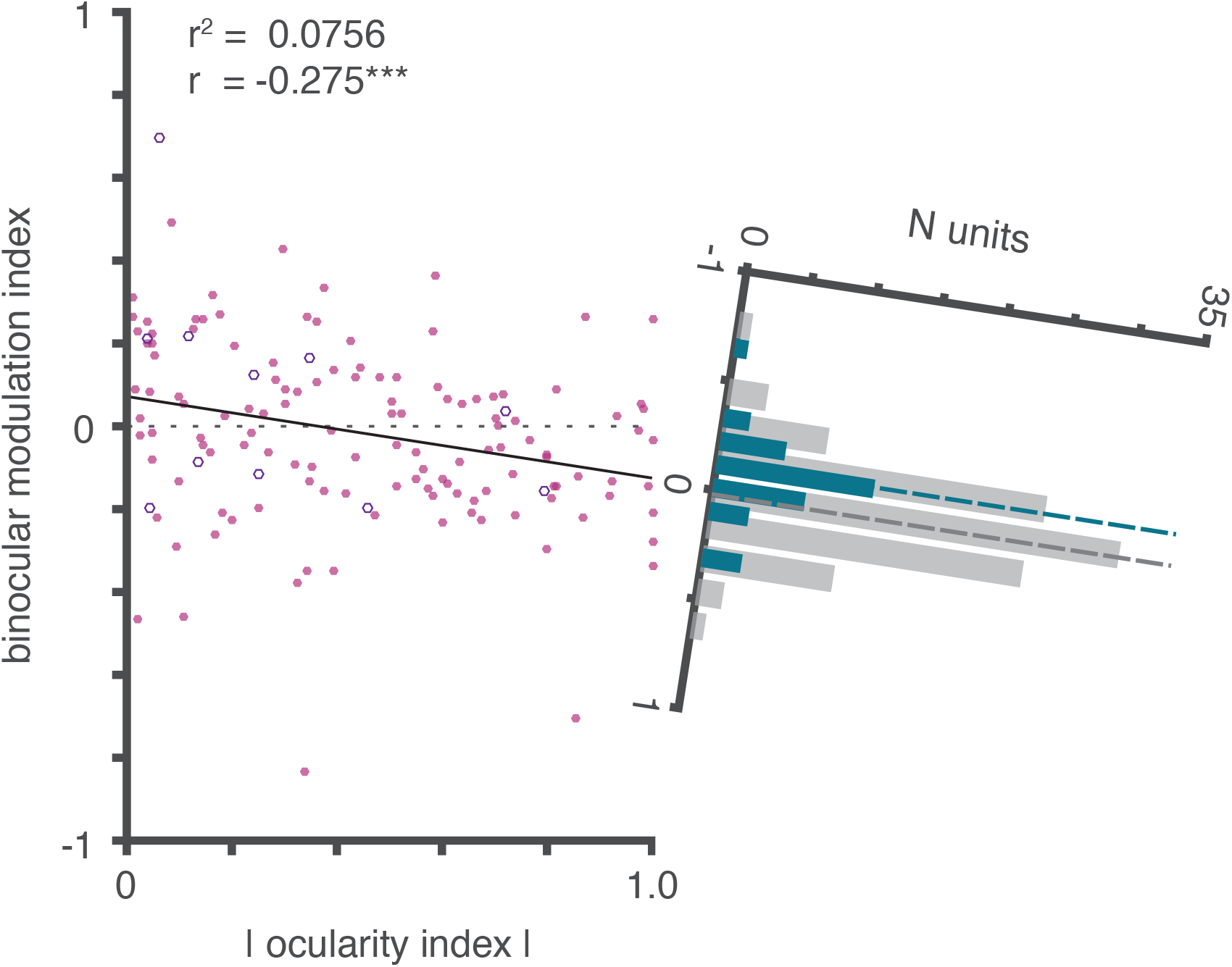
Ocular Dominance Correlates with Binocular Modulation. Rectified ocularity index versus binocular modulation index across monocular neurons (n = 138). Hollow circles represent neurons from monkey I34. Higher ocularity indices indicate more pronounced monocular responses. Lower binocular modulation indices indicate greater degree of suppression during binocular stimulation. The solid black line represents a linear regression using least-squares (p = 0.0011). The dashed line indicates the expected relationship if there were no systematic response differences between a monocular neuron’s preference for one eye and its binocular modulation. The histogram to the right shows the spread of binocular modulation for monocular (teal) and binocular (gray) neurons, respectively. The dashed vertical lines represent the median binocular modulation index for each group of neurons.

While ocular dominance explains only a small part of the overall variance of binocular modulation, this linear relationship suggests that the more a neuron prefers one eye over the other, the more binocular stimulation suppresses that neuron. We further examined this relationship as a function of laminar compartment using the multiunit population, and observed a significant effect in upper, middle, and lower layers, with a more pronounced correlation in the former two (**Figure S2**). Congruent with this correlation, responses to binocular stimulation were suppressed for most of the monocular neurons, making inhibitory interactions the predominant form of binocular interactions at the input stage of V1.

## DISCUSSION

This study is the first to our knowledge demonstrating that almost all primate V1 neurons, including those in layer 4C, are sensitive to what is shown to both eyes. Specifically, we show that monocular neurons in layer 4C, thought to receive the bulk of geniculate projections, in fact encode a binocular signal. This finding suggests that established models of binocular processing that segregate monocular signals in layer 4 (**Figure 5a**) (Hubel and Wiesel, 1972) need revision (**Figure 5b**). Moreover, we found that the more a V1 neuron responds to one eye, the more its responses are suppressed when both eyes are stimulated simultaneously. This result is significant because several impactful theoretical models on binocular vision rest on the idea that monocular neurons are inhibited by activation of the neurons’ non-dominant eye (Blake, 1989, Read et al., 2002, Ding and Sperling, 2006, Meese et al., 2006, Said and Heeger, 2013), but until now empirical evidence for this conjecture has been lacking.

**Figure 5.**
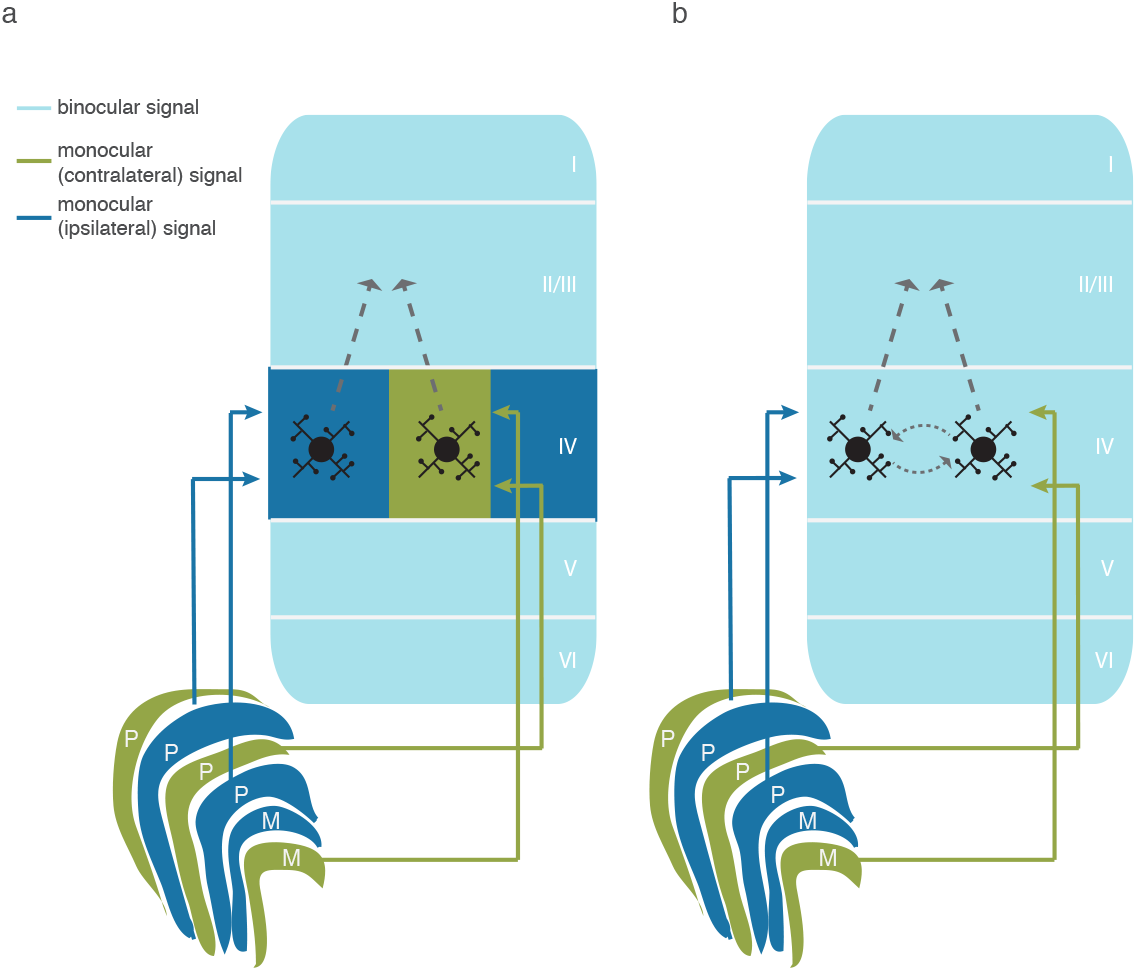
Schematic Model of Formation of Binocular Signals in V1. (a) Earlier models of binocular processing purported that monocular signals exit the LGN and remain segregated after targeting monocular neurons in L4C of V1. Binocular convergence was expected to occur outside of layer 4C. Consequently, most neurons outside layer 4C encode a binocular signal while retaining preference for one over the other eye. (b) Suggested modified model that incorporates the results from this study. While most LGN neurons are sensitive to one eye only, their target neurons in V1 layer 4C are sensitive to both eyes. Accordingly, these neurons, like those in all other layers of V1, encode what is shown to both eyes, even if they are not explicitly driven by stimuli in one of the two eyes.

### Relation to Prior Work

The findings reported here parallel similar observations in cat area 17 (Shatz et al., 1977, LeVay et al., 1978, Kato et al., 1981). However, binocular processing is implemented fundamentally differently in cats than in primates. For example, eye-specific terminations of LGN neurons in layer 4 of visual cortex are far less distinct in cats than they are in monkeys, resulting in less anatomical segregation of monocular signals in the main input layer in cats (Hubel and Wiesel, 1972, Wiesel et al., 1974, LeVay et al., 1978). Moreover, cat LGN features anatomical connections between monocular layers that are either absent or less prominent in primates (Hayhow, 1958, Laties and Sprague, 1966, Guillery and Colonnier, 1970, Sanderson et al., 1971, Saini et al., 1981). Accordingly, responses of the vast majority of cat LGN cells modulate under binocular stimulation, whereas only a small minority of LGN neurons in the macaque seem to be sensitive to both eyes (see Dougherty et al., 2018 for a more extensive discussion of species differences). Our results suggest that despite these species differences, monocular neurons in primary visual cortex of both cats and monkeys exhibit binocular modulation.

Given these analogous results, we decided to use the previously published cat data to compute a statistical power analysis. The result showed that for the reported effect size and estimated variance, a sample of 11 neurons yields 80% power, suggesting that our sample size – though small – offered sufficient degrees of freedom (Kato et al., 1981).

Previous work in primates inferred that virtually all V1 neurons are sensitive to interocular disparity (Poggio and Fischer, 1977). This idea is corroborated by our finding that virtually all V1 neurons – including monocular neurons in layer 4C – carry binocular signals. Several primate studies also quantified the fraction of primate V1 cells that respond when a stimulus is shown to one or both eyes (Kiorpes et al., 1998, Macknik and Martinez-Conde, 2004), and some considered ocularity across cortical depth (Hubel and Wiesel, 1968, Schiller et al., 1976). Several of these studies used a scale to rate the extent to which one eye or the other drives neurons (Schiller et al., 1976, Kiorpes et al., 1998, Parker, 2007). This ocular dominance scale consists of 7 distinct groups, with groups 1 and 7 corresponding to neurons driven exclusively by the contra- and ipsilateral eyes, respectively (Hubel and Wiesel, 1962, Hubel and Wiesel, 1968). Group 4 corresponds to neurons driven equally through both eyes. Using this technique, some authors reported distributions appearing Gaussian (Parker, 2007), matching our finding, while others reported more uniform distributions (Kiorpes et al., 1998). Reasons for this variance might include the subjective nature of the rating system and the laminar position of neurons sampled. Importantly, though, previous studies and the data reported here agree that only a small fraction of V1 neurons is monocular (Baker, 1974).

### Binocular Modulation and Disparity Tuning

We did not systematically test our recorded neurons for disparity tuning. In other words, we presented all of our binocular stimuli at zero disparity only. It is possible that some of the neurons in our sample that did not show a significant effect of binocular modulation might have actually been sensitive to both eyes. For example, if these neurons were disparity tuned and preferring non-zero disparity, a significant effect of binocular modulation might have been evident if we had shown the stimuli at the preferred disparity. Indeed, several previous studies in both cats and monkeys found disparity tuning among monocular V1 neurons (Ohzawa and Freeman, 1986, Prince et al., 2002, Read and Cumming, 2004). Our findings confirm and expand on these studies by demonstrating that a large fraction of monocular neurons V1, including those in layer 4C, are sensitive to both eyes even if no disparity is present, and demonstrate that even at zero disparity there is a correlation between binocular modulation and ocularity.

### Possible Explanations for the Binocular Modulation of Monocular Neurons

The binocular response modulation of monocular neurons that we observed could arise through one of several mechanisms, or a combination thereof. One possibility is that monocular neurons in layer 4C of V1 receive subthreshold inputs from their non-dominant eye. Indeed, some geniculate projections to V1 layer 4C have been shown to bifurcate and innervate neighboring ocular dominance columns (Blasdel and Lund, 1983). These axons might form—possibly less potent—connections with neurons in ocular dominance columns of the other eye, thus leading to binocular convergence at the thalamo-cortical synapse. However, this kind of connectivity could only explain binocular facilitation, and not binocular suppression, because geniculate projections to V1 are excitatory. A second possibility is that intralaminar interactions among layer 4C neurons, including those by bridging interneurons, cross ocular dominance columns. This connectivity would allow for cross-talk between the signals of each eye (Ahmed et al., 1994, Katz et al., 1989), again placing the locus of binocular convergence within layer 4C. A third possibility is that excitatory and inhibitory neurons in other layers target layer 4C neurons through interlaminar connections. These interlaminar connections might feed binocular signals back to these monocular cells in layer 4C (Gilbert and Wiesel, 1989, Wiser and Callaway, 1997). In this case, intracellular summation of monocular inputs from either eye occurs outside of layer 4C. The resulting binocular responses are then fed back to layer 4C, causing binocular modulation that is secondary to binocular convergence.

Interestingly, several empirical and psychophysical studies suggest that monocular neurons interact at or before the point where monocular signals merge into a binocular signal (Truchard et al., 2000, Ding and Sperling, 2006). Human functional magnetic resonance imaging corroborates this prediction (Moradi and Heeger, 2009). Given that, the most parsimonious interpretation of our data is that monocular neurons in layer 4C interact directly (see also below).

### Binocular Signals in The Lateral Geniculate Nucleus

Another important consideration is that binocular processing may initiate in the LGN. Specifically, LGN monocular neurons might modulate under binocular stimulation and imprint this response pattern onto their projection targets in layer 4C of V1. Indeed, binocular modulation has been reported in cat LGN (see Dougherty et al., 2018 for review). Moreover, cortical inactivation studies aimed at delineating whether this binocular modulation is caused by feedback from V1 produced equivocal results (Singer, 1970, Sanderson et al., 1971, Schmielau and Singer, 1977, Pape and Eysel, 1986, Varela and Singer, 1987). These findings leave open the possibility that LGN neurons interact on the subcortical level, resulting in binocular modulation prior to V1.

Importantly, however, the findings in the cat could not be replicated in primates, suggesting that the two species differ substantially in their functional organization of binocular integration (Rodieck and Dreher, 1979). More work is needed to determine the degree of binocular modulation in primate LGN as well as whether it is fed back from V1 or local in origin. Nonetheless, it is worth noting that a small fraction of primate LGN neurons can be driven through either eye (Cheong et al., 2013, Zeater et al., 2015). These neurons comprise a subset of the koniocellular neurons. Whether these binocular responses are supported by local neural interactions or fed back from cortex is unknown. Taken together, these observations suggest that the vast majority, if not all, of geniculate inputs to layer 4C do not encode binocular signals in primates. Therefore, the binocular modulation of monocular neurons, reported here, is likely of cortical origin.

## ACKNOWLEDGMENTS

The authors would like to thank Dr. W. Zinke, S. Amemori, Dr. T. Apple, M. Feurtado, K. George-Durrett, Dr. A. Graybiel, N. Halper, P. Henry, M. Johnson, Dr. C. Jones, M. Maddox, L. McIntosh, Dr. A. Newton, J. Parker, C. Thompson, K. Torab, C. Subraveti, B. Williams and R. Williams for technical advice and assistance. We thank Dr. B. Cumming, B.M. Carlson and N. Valov for comments on an earlier version of this manuscript. This work was supported by a research grant from the National Eye Institute (1R01EY027402-01). K.D. and J.A.W. are supported by a National Eye Institute Training Grant (5T32 EY007135-23). A.M. is supported by a research grant of the Whitehall Foundation, a Career Starter grant by the Knights Templar Eye Foundation, and a Fellowship of the Alfred P. Sloan Foundation.

## AUTHOR CONTRIBUTIONS

K.D., M.A.C., and A.M conceptualized and designed the study. K.D and M.A.C. trained animals and implemented the experiments. K.D., M.A.C. and J.A.W. collected neurophysiological data and preprocessed the data. K.D. performed the data analysis and prepared the figures for publication. K.D. and A.M. wrote the manuscript with input from other authors.

## STAR METHODS

### CONTACT FOR REAGENT AND RESOURCE SHARING

Information and requests for resources used in this study should be directed to Dr. Alexander Maier.

### EXPERIMENTAL MODEL AND SUBJECT DETAILS

Two adult monkeys (*Macaca radiata*, one female) were used in this study. Both animals were pair-housed. Both animals were on a 12-hour light-dark cycle, and all experimental procedures were carried out in the daytime. Each monkey received nutrient-rich, primate-specific food pellets twice a day, along with fresh produce and other forms of environmental enrichment at least five times a week. All procedures followed regulations by the Association for the Assessment and Accreditation of Laboratory Animal Care (AAALAC), Vanderbilt University’s Institutional Animal Care and Use Committee (IACUC) and National Institutes of Health (NIH) guidelines.

## METHOD DETAILS

### Surgical Procedures

Prior to data collection, each monkey was implanted with a custom-designed plastic head holder and a plastic recording chamber (Crist Instruments) in two separate surgeries under sterile conditions. The animals were administered isoflurane anesthesia (1.5-2.0%). Vital signs, including blood pressure, heart rate, SpO_2_, CO_2_, respiratory rate and body temperature were continuously monitored throughout the whole procedure. During surgery, the head holder or the recording chamber was attached to the skull using transcranial ceramic screws (Thomas Recording) and self-curing dental acrylic (Lang Dental Manufacturing). A craniotomy was performed over the perifoveal visual field representation of primary visual cortex (V1) in each monkey concurrent with the positioning of the recording chamber. Each monkey was given analgesics and antibiotics, and closely observed by researchers, facility veterinarians and animal care staff for at least three days following surgery.

### Data Acquisition and Pre-Processing

During each recording session, a linear multielectrode array (U-Probe, Plexon Inc., or Vector Array, NeuroNexus) with either 24 or 32 contacts of 0.1 mm inter-contact spacing was carefully inserted into V1. Extracellular voltage fluctuations (0.5 Hz – 30 kHz) were recorded inside an electromagnetic radio frequency-shielded booth. These signals were amplified, filtered and digitized using a 128-channel Cerebus^®^ Neural Signal Processing System (NSP; Blackrock Microsystems). Both a broadband (0.3 Hz – 7.5 kHz) signal sampled at 30 kHz and a low frequency-dominated signal (0.3 Hz – 500 Hz) sampled at 1 kHz was saved for offline analysis. The NSP also recorded the output of a photodiode signal (OSI Optoelectronics) placed on the monitor to track stimulus-related events at 30 kHz. The NSP further digitized the output of the optical eye tracking system (EyeLink II, SR Research or SensoMotoric Instruments) at 1 kHz, as well as digital event markers sent from the behavioral control system (MonkeyLogic, Asaad et al., 2013). Both the photodiode signal and event markers were used to align the neural data with visual and behavioral events.

All neurophysiological signals, except for local field potentials (LFP), were extracted offline from the digitized broadband signal using custom written code in MATLAB (2016a; The Mathworks, Inc.). LFP was extracted from the low frequency-dominated signal described above.

We extracted multiunits by applying a time-varying threshold to the envelope of the broadband signal, and saved all time points where the signal envelope exceeded a preset threshold. Specifically, we first lowpass-filtered the 30 kHz-sampled voltage signal at 5 kHz using a second order Butterworth filter. We then downsampled the signal by a factor of 3. Next, we high pass-filtered the signal at 1 kHz cut-off with a second-order Butterworth filter. Finally, we rectified the resulting data. To compute the signal envelope, we downsampled the signal by a factor of 3. To compute a threshold, we smoothed the signal by convolving the data with a 1 s boxcar function and then multiplied the result by 2.2. To recover temporal information, we extracted +/− 0.3 ms of data from the original signal for each time point where the envelope exceeded the threshold. We then adjusted these time points to correspond to the point of maximum slope within this window.

For laminar alignment (see below), we used an analog multiunit signal that was computed by high-pass filtering the broadband signal at 750 Hz with a fourth-order Butterworth filter, followed by a full-wave rectification step.

We extracted single neurons with KiloSort, an open-source unsupervised machine-learning algorithm for spike-sorting (Pachitariu et al., 2016), using the default parameters for sorting and cluster merging. We extracted +/− 1 ms of data around each KiloSort’ed spike time from the original broadband signal for each simultaneously recorded electrode contact. We then averaged across impulses to create a spatiotemporal map of the spike waveform (time x electrode contacts). The region of the spatiotemporal waveform map that exceeded +/− 30% of maximum modulus had to span fewer than 3 electrode contacts (0.3 mm) and 0.9 ms to be included in the study. Neurons that met these criteria were localized to the electrode contact where they evoked the largest amplitude.

Spike rates were downsampled to 1 kHz. For each neuron, spike times were converted to a time-varying signal (spike density function) using 0 to represent time points without a spike and 1 for time points where a spike was detected. This time-varying signal was then convolved using a Poisson distribution resembling a postsynaptic potential (Sayer et al., 1990), with the spike rate (*R*) computed at time (*t*):

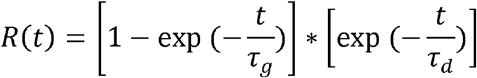

where *τ_g_* and *τ_d_* are the time constants for growth and decay, respectively. Values of 1 and 20 for *τ_g_* and *τ_d_* respectively were used based on a previous study (Hanes et al., 1995). After convolution, the signal was multiplied by the sampling frequency to convert units to spikes per second.

Current source density (CSD) analysis was performed on the LFP signal using an estimate of the second spatial derivative appropriate for electrodes with multiple contact points (Nicholson and Freeman, 1975):

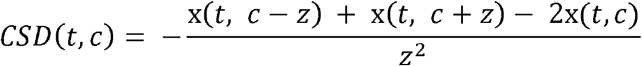

where *x* is the extracellular voltage recorded in Volts at time *t* from an electrode contact at position *c*, and *z* is the electrode inter-contact distance (0.1 mm). In order to yield CSD in units of current per unit volume, the resulting CSD from the formula above was multiplied by 0.4 S/mm as an estimate of cortical conductivity (Logothetis et al., 2007).

Eye position was measured continuously using a commercially eye tracker (see details below). Using the horizontal and vertical gaze position data provided by this system, we extracted microsaccades using a previously published algorithm. (Otero-Millan et al., 2014).

### Visual Display

Stimuli were presented on a linearized CRT monitor with a refresh rate of either 60 Hz (resolution 1280 × 1024) or 85 Hz (resolution 1024 × 768). These visual stimuli were generated using custom-written code for MonkeyLogic (Asaad et al., 2013) in MATLAB (R2012-2014, The MathWorks) on a PC (Dell, Windows 7 or Windows 10) with a NVIDIA graphics card. Animals viewed all stimuli through a custom-built mirror stereoscope that employed infrared-light passing cold mirrors (Edmund Optics). The animal, mirrors and monitor were positioned so that the animal’s right eye viewed stimuli presented on the right side of the monitor and the animal’s left eye viewed stimuli on the left side of the monitor. To prevent light scatter from one side of the monitor to the opposing eye, a black, non-reflective septum was placed between the monitor and the back side of the mirrors, effectively dividing the left and right sides of the apparatus.

Infrared-light sensitive cameras, placed directly behind the cold mirrors on the stereoscope, were used to track gaze position with commercially available eye tracking software (Eye Link II, SR Research). Gaze position was converted to an analog signal and inputted to MonkeyLogic/MATLAB (NIDAQ PCI-6229) at 1 kHz. At the beginning of each recording session, the stereoscope was calibrated to facilitate binocular fusion of the left and right sides of the monitor using a behavioral task that relied on acquiring the same gaze position for corresponding locations on each side of the monitor (Maier et al., 2007, Maier et al., 2008). To further aid fusion, an oval aperture or set of intersecting circles in each corner was displayed at the edge of each half-screen.

### Laminar Alignment and RF Mapping

For each penetration with the linear multielectrode array, CSD analysis was used to locate the boundary between layer 4C and layer 5. CSD analysis of visual responses to brief visual stimulation has been shown to reliably indicate the location of the primary geniculate input to V1 (in granular layer 4C, or L4) by a distinct current sink that is thought to reflect the combined excitatory post-synaptic potentials of the initial retino-geniculate volley of activation (Mitzdorf and Singer, 1977, Schroeder et al., 1998). Analog multiunit responses, or more precisely lack thereof, were used to identify electrode contacts that lie outside V1, either in the subdural space or the white matter below. We excluded contacts on the extreme ends of the array that did not exhibit a visual response. After removing these contacts, the location of the initial current sink was used to align and average data across electrode penetrations, resulting in 0.1 mm +/− 0.05 mm resolution across the depth of V1 (Maier et al., 2010, Maier et al., 2011, Godlove et al., 2014, van Kerkoerle et al., 2014, Hansen et al., 2012, Spaak et al., 2012, Ninomiya et al., 2015, Cox et al., 2017, Dougherty et al., 2017).

For display, representations of CSD as a function of time and space were Gaussian-filtered (σ = 0.1). Electrode contacts were classified to be in supragranular, granular, or infragranular positions based on the CSD responses as well as neurophysiological criteria. These criteria included the power spectral density of the LFP across cortical depth, signal correlations of the LFP between all contact combinations, and stimulus-evoked analog multiunit responses. The supragranular-to-granular boundary is more challenging to define based on these criteria and was instead set to 0.5 mm above the granular to infragranular boundary.

Once the linear multielectrode array was appropriately positioned in cortex, a reverse correlation-like technique was used to map the RFs of the neurons under study. In each trial, the animals fixated while up to five circular Gabor-filtered static random noise patches appeared in sequence at pseudorandom locations within a pre-defined virtual grid of monitor locations. Each noise patch was displayed for 150 ms with an inter-stimulus interval of 150 ms. The size of each noise patch and the size of the pre-defined grid depended on the recording session. Typically, each session included a “coarse” mapping phase to determine the general location of the RF. We then used a subsequent “fine” mapping phase to map the precise location of the RF. We then used 3D Receptive Field Matrices (RFMs) (Cox et al., 2013) to create a map of each electrode contact’s neuronal response for different points in visual space (see **Figure S4a**). For every multiunit or single neuron, we averaged the spiking response to each stimulus presentation across time, resulting in a single scalar value. We then converted these scalar values to units of z-score. We filled the retinotopic portion of the RFM corresponding to the stimulus location with the z-score for every presentation. This procedure produced a 3D matrix, with two dimensions representing vertical and horizontal visual space and a third dimension representing the response magnitude for each multiunit or neuron. We then averaged this third dimension to produce a spatial map of the mean response. We computed RF centers and extents by fitting an oval to the largest, contiguous patch of the spatial map that exceeded 1 z-score.

### Monocular and Binocular Visual Stimulation

We trained each animal trained to fixate on a small (0.2 degrees of visual angle, dva) cross presented at the center of each eye’s visual field. Animals held fixation for several (< 5) seconds while we presented stimuli in their perifoveal visual field. The results reported in this paper are based on data that come from three different paradigms, all with the same or similar conditions. For all neurons, we recorded responses to high contrast (0.8 or greater) sine-wave gratings at varied orientations presented to the left eye or right eye. For 138 neurons, we also recorded responses to high contrast sine-wave gratings at varied orientations presented to both eyes simultaneously. For fewer neurons (see **Table 2, Table S1**), we recorded responses to sine-wave gratings where the contrast in the two eyes varied at several different levels. We presented all stimuli for at least 200 ms, and limited the data to the initial 200 ms of stimulus presentation for each neuron. Where data for multiple paradigms existed for one neuron, we concatenated data across the same conditions.

**Table 2.** Number of neurons for computing the contrast response functions shown in Figure 2.

|  | dominant eye<br>Michelson<br>contrast | 0.0 | 0.2 – 0.3 | 0.45 – 0.5 | 0.8 – 1.0 |
| --- | --- | --- | --- | --- | --- |
| suppressed<br>group | monocular<br>conditions | n/a | 16 | 16 | 23 |
|  | binocular<br>conditions | 23 | 15 | 16 | 23 |
| facilitated<br>group | monocular<br>conditions | n/a | 5 | 6 | 10 |
|  | binocular<br>conditions | 10 | 4 | 4 | 10 |

In one paradigm, animals fixated on a fixation cross for at least 300 ms before a sequence of up to five circular sinusoidal gratings appeared. Each grating was presented for at least 200 ms (46 sessions) or 500 ms (23 sessions) before an inter-stimulus interval of at least 200 ms. We presented the gratings randomly to either the left eye, right eye, or both eyes over the population RF location of the recorded neurons. The grating stimuli varied in orientation but always had a Michelson contrast above 0.8 (mode: 0.9) as well as constant spatial frequency (0.5-3 cycles/deg). If the animals successfully held fixation within a 1 dva radius around the fixation cross for the entire stimulus sequence, liquid juice reward was delivered. If the animals broke fixation or blinked, the trial was aborted and a short timeout (1-5 s) was given before the start of the next trial.

In the second paradigm, we used the same parameters, including stimulus timing. However, we presented the gratings at only one of two orientations (the neurons’ estimated preferred orientation or the orientation orthogonal to this preferred orientation) and varied contrast of the gratings shown to each eye across trials (see **Table 2, Table S1**). We determined the preferred orientation based on online analyses of the multiunit responses to sine-wave gratings of varying orientations. If preferred orientation varied across electrode contacts, we chose the preferred orientation shared by the most number of contacts.

In the third paradigm, the animals fixated for at least 300 ms before we presented gratings at the same orientation in both eyes (either the preferred or non-preferred orientation, as described for paradigm two) and varied contrast of the gratings shown to each eye across trials (see **Table 2, Table S1**). Stimuli were shown for 1600 ms (12 sessions). If the animals successfully held fixation within a 1 dva radius around the fixation cross for the entire stimulus duration, liquid juice reward was delivered. If the animals broke fixation or blinked, the trial was aborted and a short timeout (1-5 s) was given before the start of the next trial.

## MRI

Animals were anesthetized using the same procedure as outlined under *Animal Care and Surgical Procedures*. Anesthetized animals were placed inside a Philips Achieva 7T MRI scanner at the Vanderbilt University Institute of Imaging Science and remained anesthetized throughout the duration of the scan. Vital signs were monitored continuously. T1-weighted 3D MPRAGE scans were acquired with a 32-channel head coil equipped for SENSE imaging. Images were acquired using a 0.5 mm isotropic voxel resolution with the following parameters: repetition time (TR) 5 s, echo time (TE) 2.5 ms, flip angle 7°.

## QUANTIFICATION AND STATISTICAL ANALYSIS

For a KiloSort’ed neuron or multiunit to be considered for analysis, it had to be located within the grey matter (see **Laminar Alignment and RF mapping**). Moreover, the neuron or multiunit’s mean initial response (40-100 ms) to the dominant eye (defined as the eye that yielded the highest mean spike rate between 40 ms and 140 ms when stimulated with a high contrast stimulus) had to exceed a maximum of 10 spikes per second. This response also had to be significantly larger than the fixation baseline (baseline window: −50-0 ms, paired t-test, p < 0.05). Lastly, there had to be at least 12 successfully completed presentations of each contralateral and ipsilateral eye stimulation using the high contrast gratings.

To compute normalized spiking, we transformed the mean responses for each neuron to z-scores. Specifically, we first subtracted their baseline firing rate. Then, we divided this value by the difference between the maximum firing rate to stimulation of the dominant eye and the baseline firing rate. Similarly, we normalized contrast response data across conditions for each neuron by subtracting the baseline firing rate from the mean response at each contrast level.

Then, we divided each resulting value by the difference between high contrast dominant eye stimulation and baseline firing.

All statistical hypothesis tests, including Wilcoxon signed-rank tests (one and two-sided), ANOVAs with post-hoc Tukey HSD tests, Chi-square goodness of fit tests, and Pearson’s correlation analysis, are fully described where used. All reported confidence intervals were based on bootstrapping using 10,000 repetitions on the group statistic (mean or median) shown.

Neurons were included in the monocular category if they had a non-significant dominant eye response during the initial stimulation window (40 to 100 ms) relative to baseline (−50 to 0 ms) (one-tailed t-test, p < 0.05). We re-categorized two neurons as binocular following visual inspection of average responses.

To quantify the relative amount of excitation by stimulation of the contralateral eye versus by that of the ipsilateral eye, we calculated an ocularity index for each unit:

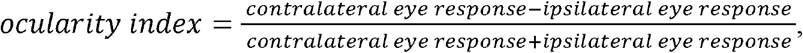

where response was defined as the half-wave rectified, baseline-subtracted mean spike rate during the initial response period (40-140 ms). We calculated a binocular modulation index to assess the strength and direction of binocular modulation using the following formula:

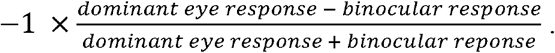

We calculated response latency for each neuron using a custom algorithm. Briefly, we used the z-scored response to stimulation in the dominant eye to determine the first time point that exceeded a threshold while trending in positive direction. Specifically, we parceled the data into overlapping windows, whose length was defined as 3% of the maximum response time. We then defined a threshold as the mean response plus two standard deviations for the time between 15 ms before to 15 ms after stimulus onset. If the resulting threshold was lower than 0.05, it was set to 0.05. Criterion was met if 90% of data points within a window exceeded this threshold while 70% of data points trended positively. If no data point fit those criteria, we used the first time point that crossed threshold instead.

In addition to group statistics, we used receiver-operating characteristics (ROC) analysis (Green and Swets, 1966) to determine whether there was a significant difference between stimulation conditions at the single-neuron level. Specifically, for each neuron, we ran an ROC analysis with twelve thresholds using 10 ms bins of data, and a sliding window of 1 ms during the response period (20-190 ms). Statistical significance was determined by comparing the area under the curve to a bootstrapped distribution of area under the curve values computed on 10,000 repetitions of shuffled data.

## DATA AND SOFTWARE AVAILABILITY

Code used for analyses in this paper are available upon request from the corresponding author.

